# Machine learning Classification of Dyslexic Children based on EEG Local Network Features

**DOI:** 10.1101/569996

**Authors:** Z. Rezvani, M. Zare, G. Žarić, M. Bonte, J. Tijms, M.W. Van der Molen, G. Fraga González

## Abstract

Machine learning can be used to find meaningful patterns characterizing individual differences. Deploying a machine learning classifier fed by local features derived from graph analysis of electroencephalographic (EEG) data, we aimed at designing a neurobiologically-based classifier to differentiate two groups of children, one group with and the other group without dyslexia, in a robust way. We used EEG resting-state data of 29 dyslexics and 15 typical readers in grade 3, and calculated weighted connectivity matrices for multiple frequency bands using the phase lag index (PLI). From the connectivity matrices, we derived weighted connectivity graphs. A number of local network measures were computed from those graphs, and 37 False Discovery Rate (FDR) corrected features were selected as input to a Support Vector Machine (SVM) and a common *K* Nearest Neighbors (KNN) classifier. Cross validation was employed to assess the machine-learning performance and random shuffling to assure the performance appropriateness of the classifier and avoid features overfitting. The best performance was for the SVM using a polynomial kernel. Children were classified with 95% accuracy based on local network features from different frequency bands. The automatic classification techniques applied to EEG graph measures showed to be both robust and reliable in distinguishing between typical and dyslexic readers.

## 1. Introduction

Developmental dyslexia is a specific reading and spelling disability with a genetic and neurobiological component and relatively high prevalence rates around 5% (Blomert 2005; Snowling 2013). Neuroimaging studies have investigated biomarkers of dyslexia using structural and functional network analyses and there is a growing interest in connectivity abnormalities between different brain systems that may result in impaired reading (e.g., Finn et al. 2014; Schurz et al. 2014; Liu et al. 2015). Diffusion tensor imaging studies show reduced connectivity in the main white matter pathways (see review in Vandermosten et al. 2012). Functional magnetic resonance imaging (fMRI) studies reported reduced long-range connectivity across brain systems specialized for reading (e.g., Pugh et al. 2000; Shaywitz et al. 2003; van der Mark et al. 2011; Schurz et al. 2014), and associated functional networks (Wolf et al. 2010; Finn et al. 2014). Whole brain connectivity studies observed that dyslexics differ from typical readers in whole brain networks organization, showing increased local processing and less long-range communication (Liu et al. 2015) and lower global efficiency in dyslexics (Dimitriadis et al. 2018). Collectively, this evidence supports the view that a widespread network of brain regions may be compromised in developmental dyslexia (Martin et al. 2016).

An important objective of the neuroscientific search for biomarkers of dyslexia is to contribute to the diagnosis and early detection or prediction of reading disabilities. The classification of individuals suffering from several neuropsychological disorders may take advantage of using machine learning methods (e.g., Duda et al. 2016; Kessler et al. 2016). These methods are particularly useful when applied to neuroimaging data as they allow for using widely distributed information to improve the classification of clinical groups or individuals at risk. A recent MRI study used a machine learning classifier and reported above chance levels in discriminating dyslexic adults from controls based on gray matter differences (Tamboer et al. 2016). Two related studies employed multivariate pattern analysis of brain activity during a phonological task to identify poor readers (Tanaka et al. 2011) and predict long-term outcomes in dyslexic children based on whole-brain activation (Hoeft et al. 2011). The latter study showed that methods using brain measures outperformed procedures relying on behavioral measures in predicting reading improvements across the 2.5 years following the experiment. These studies illustrate the potential of using machine learning techniques in combination with neuroimaging data to improve the classification and early detection of dyslexia.

The present study uses a ML classifier to discriminate between dyslexic and typically reading children based on functional connectivity using measures derived from the electroencephalogram (EEG). It has been demonstrated previously that task-independent EEG activity contains information about how different brain systems communicate and how functional networks may be intrinsically organized (van den Heuvel and Hulshoff Pol 2010). Importantly, earlier studies related neural activity at rest to reading ability in children and adults, showing that a resting-state paradigm can be profitably used to study language networks (Hampson et al. 2006; Koyama et al. 2010).

Given the highly interactive and complex nature of reading, the study of its neurobiology might benefit from an integrative and holistic view of brain function conceptualized as a complex network (Bullmore and Sporns 2009). Within that framework, graph theoretical analysis allows for modeling whole-brain functional connectivity networks as a set of nodes (vertices) and the connections between them (edges). The multiple measures that can be derived from a graph are used to describe the network in terms of information transfer and balance between ‘segregation’ and ‘integration’ (see reviews in Bullmore and Sporns 2009, 2012). Two magnetoencephalographic (MEG) studies of dyslexia examined graph measures and found dysfunctional long-and short-range functional connectivity in dyslexics during a reading task (Vourkas et al. 2011) and less organized connectivity at rest (Dimitriadis et al. 2013). In a previous study, we applied graph analysis to resting-state EEG to compare dyslexics and typically reading children in grade 3 (Fraga González et al. 2016). The results suggested group differences in several *global* graph metrics in the theta band suggesting a reduced network integration and communication between the nodes in dyslexics compared to typical readers.

In our previous graph study, we examined global properties of the network that were described by graph measures averaged across electrodes (Fraga González et al. 2016). For the current analysis, we will employ data from that study, however, we use a different analysis approach by extracting local features, i.e., computed per node, and take a step further to clinical application and deploy artificial intelligence. This local information could reflect aspects of regional connectivity organization relevant to network development in dyslexia (Liu et al. 2015) and may provide neural features for the benefit of the SVM classifier performance. Basically, an SVM is a discriminative classifier formally defined by a hyperplane. Given labeled training data (supervised learning), the algorithm outputs an optimal hyperplane that categorizes new examples. Due to its ability to manage large datasets, the algorithm is widely used for binary classification problems in machine learning. For more details on SVM see (Hsu et al. 2003). We will use methodological approach that is similar to the one applied in a previous study that, using resting-state EEG and an SVM classifier, identified 6-month-old infants at familial risk for a language learning disorder (Zare et al. 2016). The current study uses SVM and KNN to classify children in 3^rd^ grade as dyslexics or typical readers, based upon a large number of local features derived from functional connectivity matrices in the different frequency bands of the EEG. Cross-validation is employed to assess the resulting classification, and random shuffling is deployed to assure that the classifying performance is not due to bias in feature selection criteria. We aimed at assessing the utility of machine learning techniques to find best distinguishable characteristics in reading difficulties based on functional EEG networks. To the best of our knowledge, this is the first study to use local EEG features to designing a classifier for dyslexia.

## 2. Methodology

The current analysis is performed on data from a previous study (Fraga González et al. 2016). We refer to that article for a more extensive description of the EEG recordings and summarize here only the information needed for the current study.

### 2.1. Participants

The participants of the current study were 44 subjects including 29 third-grade dyslexic children (mean age = 8.96; SD = 0.40); with a percentile score of 10 or lower on a standard reading test, and 15 third-grade children (8.75 ± 0.31 years old) in the control group with the same socio demographical background as dyslexic; with no history of reading difficulties and had a percentile score of 25 or higher on standard reading tests. The participants of the current study were part of a larger sample of 62 children participating in a larger study. Due to young age of our participants some participants were excluded due to excessive movement or other artifacts in the data, or did not complete the resting-state recordings. All participants were native Dutch speakers, received two and a half years of formal reading instruction in primary education. The study was approved by the Ethical Review Board of the University and all parents or caretakers signed informed consent before the children participated. Demographic characteristics and reading scores of the complete sample are included in S1 Table.

### 2.2. EEG recording and preprocessing

The total duration of the eyes-closed resting state data collected was 2 minutes. EEG data were collected using 64 channels with sampling frequency of 250 Hz; Biosemi ActiveTwo system. Data was imported in Brain Vision Analyzer (Version 2.01.5528 © Brain Products) where spline interpolation was applied to channels with excessive artifacts and segmented in 30 epochs of 4 seconds. Epochs containing excessive noise and artifacts were visually inspected and removed. For each subject, 10 artifact-free epochs were selected and exported to ASCII files. The data were imported to Brainwave v0.9.117 (developed by Prof. Cornelis Stam’ s research group; freely available at http://home.kpn.nl/stam7883/brainwave.html) where it was re-referenced to the common average, submitted to power spectral analysis using Fast Fourier Transform (FFT) and filtered into four frequency bands [delta (0.5-4Hz); theta (4-8Hz); alpha (8-13Hz); beta (13-30Hz)]. Then, the functional connectivity matrices were obtained in Brainwave and based upon those matrices weighted network measures were computed using custom MATLAB code (R2016; The Mathworks, Natick, MA), as well as with functions available as part of the MATLAB Machine learning and Statistics Toolboxes and the Brain Connectivity Toolbox (BCT) (Rubinov and Sporns 2010).

## 3. EEG analysis

### 3.1. Functional connectivity

Functional connectivity was assessed for each segment (10 segments of 4096 data points per subject) and frequency band with the phase lag index (PLI). The PLI measures the asymmetry of the distribution of phase differences between two signals. PLI provides a measure of statistical interdependencies between time series, which reflects the strength of coupling. The major aim of using the phase lag index is to obtain reliable estimates of phase synchronization that are invariant of the presence of common sources, a feature that may be absent in other connectivity measures (Stam et al. 2007). Asymmetry of the phase difference distribution indicates that the likelihood of phase difference Δ*φ* being in the interval −*π* < Δ*φ* < 0 is different from the likelihood of being in the interval 0 < Δ*φ* < *π*. This asymmetry indicates a phase difference (or ‘lag’) between the two time series (Stam et al. 2007). The adjacency matrix is constructed using formula

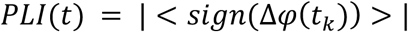

where Δ*φ* is the phase difference and *t_k_* is the *k^th^* sequence. PLI ranges between 0 and 1 where zero indicates no coupling. Angle between each pair of time series is calculated using Hilbert transform.

### 3.2. Weighted network and local feature selection

From the previous step, weighted connectivity matrices were derived for each subject. For each of the individual 64 by 64 matrices, we calculated several local features from weighted network measures per node. In a weighted graph, each electrode represents a node and all the nodes are connected by links with a specific weight representing the strength of connectivity (Barrat et al. 2004). The network measures used in this study are summarized in Table 1. We focus on the main network features that are commonly used in the literature and are measured at local level in the network (Bullmore and Sporns 2009; Rubinov and Sporns 2010. The chosen measures aim to quantify how a node contributes in the information flow in the network, in the current study we try to exploit these local characteristics for classification. The features derived from the weighted network were averaged across segments for every subject. Note that for each subject, we repeated the calculations for each connectivity matrices per frequency band and segment. For all network features, the significance of FDR (false discovery rate; Benjamini and Hochberg 1995) corrected group differences (*p* < 0.05) were examined with t-tests defined as (*t*(*F_TYP_*, *F_DYS_*), *p*-value), where *F* stands for feature name, and then FDR corrected. Then, to demonstrate that this method worked well in feature selection, we employed a random shuffling technique. These steps are described in Fig 1.

**Table 1.** Network metrics and their definitions.

| Network measure | Definition |
| --- | --- |
| Characteristic path length | The number of edges mediating between two nodes |
| Degree | The number of edges incident to a node or sum of weights of incident edges (flow) |
| Efficiency | Quantifies a network's resistance to failure on a small scale |
| Clustering coefficient | Quantifies how close a given node and its neighbors are to being a clique |
| Modularity | Fraction of the edges that fall within the given groups minus the expected fraction if edges were distributed at random |
| Eccentricity | The maximum distance between a reference's node and node $i$ of a graph |
| Betweenness centrality (BC) | The number of shortest paths running through a node $i$ |

**Fig 1.**
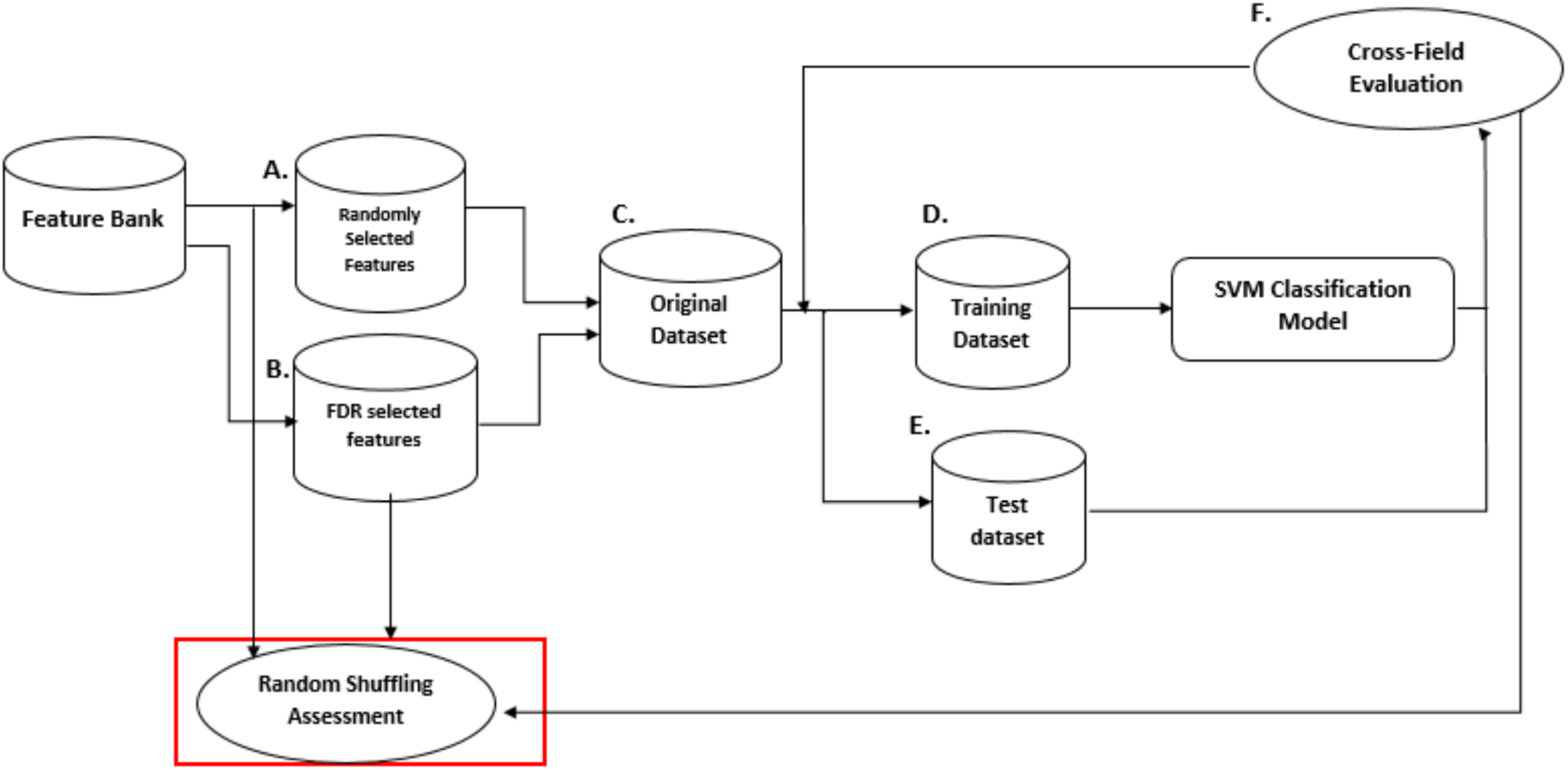
SVM classification and performance assessment by Random Shuffling. We followed two approaches to select features from those available: random selection (A) and selection via *t-*tests (B). In both cases the dataset was then divided into a Training set (D) and a Test set (E) using cross-validation. We assessed each selected feature with the SVM classifier. Finally, a random shuffling cross-fold evaluation (F) was performed to ensure the *t-*test selected features were the most relevant for classification.

#### SVM classification and cross-validation

The primary goal of this study was to use local network features for classifying children into two separate groups. In the current study, 37 features passed the FDR corrected group comparisons. Those features were used to train the classifier (see Results). Usually, it is difficult to determine in advance which classifier, and in particular which kernel function, fits the SVM classifier best from simpler to more complexity degree for a particular set of data. Therefore, starting from non-parametric classifiers to parametric classifiers, we tested several other classifiers. Here we report the classification using two kernel functions that resulted in good classification performance: linear and polynomial of degree 3. In addition, we compared the SVM to a simpler common classifier, i.e., k-nearest neighbors algorithm (KNN) with *k* = 3 and *k* = 7. In KNN classification, an object is classified by a majority vote of its neighbors, with the object being assigned to the class most common among its *k* nearest neighbors.

Subsequently, the leave-one-out-cross-validation (LOOCV) was deployed to assess the performance of the classifiers averaging performance across N sets (see Fig 1). LOOCV is the most common procedure for cross-validation where the number of sets equals the number of instances in the data set. This approach is used to avoid overfitting and promotes the reliability and generalizability of the results to a new set of data. The selected features were used as input to an SVM that performed supervised classification, mapping children into two groups: dyslexic children (DYS) and typical readers (TYP).

We assessed the classifier’s performance using the conventional measures of precision, specificity, sensitivity, and accuracy. The dyslexic reader group is designated positive and the typical readers are categorized as negative. The correct detection or “true classification” of dyslexia is then a true positive (TP). Likewise, correct classification of the typical readers is true negative (TN). Precision or Positive predictive value (PPV) indicates the proportion of positives, which are correctly identified as such (e.g., the percentage of children who are correctly identified as Dyslexic readers). Specificity refers to the percentage of participants correctly classified as typical readers, which is also known as the true negative rate (TNR). These performance indices are further described in (Zare et al. 2016).

The computations related to the SVM classifier were carried out in MATLAB 10.8.0 (R2016; The Mathworks, Natick, MA), as well as with functions available as part of the MATLAB Machine Learning Toolboxes, and in-house MATLAB code.

## 4. Results

### 4.1. PLI and network features extraction

The PLI weighted matrices that were used to extract the local features presented in Table 1. We extracted a total of 1792 features (4 frequency bands × 7 network measures × 64 channels) for each child, which were used to perform the group comparisons. Applying FDR correction, significant differences (*p* < 0.05) were obtained for a total of 37 features. The results, per frequency band, separately, are shown in Table 2. In order to visualize our results, we show an EEG channel scalp map with a color map indicating the electrode sites for which group differences were found in any of the features per frequency band (see Fig 2).

**Table 2.** FDR corrected features from weighted matrices derived from PLI.

**4.1. PLI and network features extraction**
|  |  |  |  |  |  |
| --- | --- | --- | --- | --- | --- |
| 9 |  |  | 'C6' | 0.0173 | -2.4807 |
| 10 |  | Modularity | 'AF3' | 0.0084 | -2.7708 |
| 11 |  | Efficiency | 'FPZ' | 0.0157 | -2.5194 |
| 12 |  |  | 'FP2' | 0.0015 | -3.3961 |
| 13 |  |  | 'AF8' | 0.0063 | -2.8815 |
| 14 |  |  | 'F8' | 0.0100 | -2.7027 |
| 15 |  |  | 'C6' | 0.0182 | -2.4603 |
| 16 |  | Eccentricity | 'FP1' | 0.0092 | -2.7351 |
| 17 |  |  | 'FC3' | 0.0152 | -2.5350 |
| 18 |  |  | 'CP1' | 0.0088 | -2.7497 |
| 19 |  |  | 'AF8' | 0.0000 | -4.8302 |
| 20 |  |  | 'FT8' | 0.0081 | -2.7834 |
| 21 |  |  | 'FC6' | 0.0123 | -2.6207 |
| 22 |  |  | 'C6' | 0.0072 | -2.8285 |
| 23 |  |  | 'CP6' | 0.0137 | -2.5756 |
| 24 | Theta | Betweenness centrality | 'AFZ' | 0.0014 | -3.4276 |
| 25 |  | Modularity | 'F7' | 0.0059 | 2.9055 |
| 26 |  |  | 'FT7' | 0.0049 | 2.9740 |
| 27 |  |  | 'CP3' | 0.0142 | -2.5610 |
| 28 |  |  | 'CP4' | 0.0089 | 2.7466 |
| 29 | Alpha | Betweenness centrality | 'FZ' | 0.0037 | -3.0773 |
| 30 |  |  | 'PO8' | 0.0047 | -2.9881 |
| 31 |  | Modularity | 'FC5' | 0.0197 | -2.4273 |
| 32 |  |  | 'FC3' | 0.0051 | -2.9589 |
| 33 |  |  | 'CP3' | 0.0112 | -2.6577 |
| 34 |  |  | 'PZ' | 0.0161 | -2.5099 |
| 35 |  |  | 'CPZ' | 0.0014 | -3.4211 |
| 36 |  |  | 'FC6' | 0.0060 | 2.8975 |
| 37 | Beta | Modularity | 'F4' | 0.0152 | -2.5345 |

**Fig 2.**
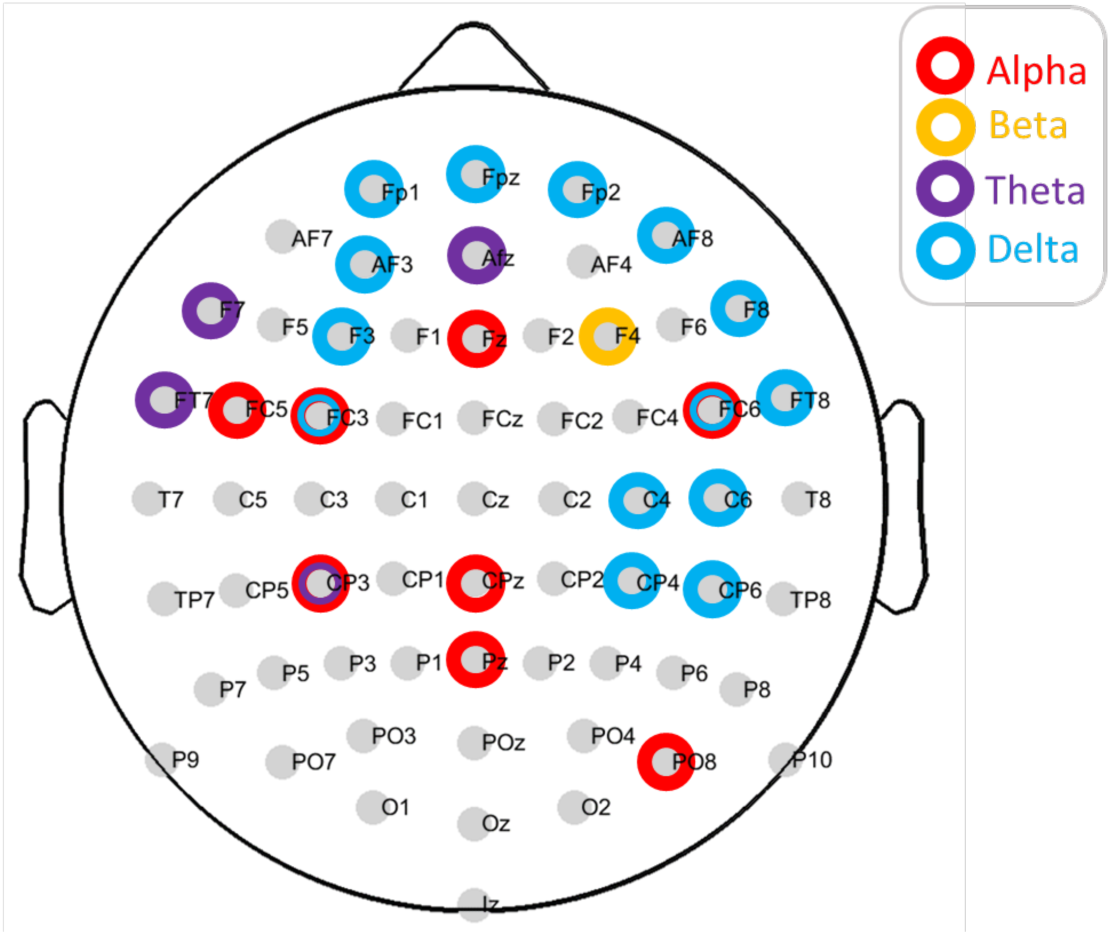
Location on scalp of EEG channels. Highlights indicate the channels where FDR corrected features were selected. The color map indicates frequency band.

#### Machine Performance

The results of machine performance after applying LOOCV are indicated in Table 3. The table shows that an SVM with a linear kernel provides the statistically best performance among the classifiers used to identify children as typical readers and dyslexic readers with high specificity, sensitivity, precision, and accuracy.

**Table 3.** Machine performance after LOOCV. Complexity of the model is reduced from left to right.

| Classifier | SVM Polynomial kernel | SVM Linear kernel | KNN, K=7 | KNN, K=3 |
| --- | --- | --- | --- | --- |
| Accuracy | 90.69 | **95.34** | 86.04 | 81.39 |
| Sensitivity | 90.00 | 96.42 | 86.66 | 85.71 |
| Specificity | 92.30 | 93.33 | 84.61 | 73.33 |
| Precision | 96.42 | 96.42 | 92.85 | 85.71 |

As input to the machine, we used a total of 37 features vectors, i.e., approximately 2% of the features, based in the FDR corrected group comparison. In order to ensure that the subset of selected features is suitable and unbiased, we used a random-shuffling method (Zare et al. 2016). In this method: a) all network features enter the classifier’s pool irrespective of the feature selection criteria. Among all network features, 37 features are randomly chosen to feed the classifier. This way, we assure that feature selection is not biased by the selection method (i.e., significant difference across two groups over a network feature); b), machine performance is evaluated using the conventional measures of precision, specificity, sensitivity, and accuracy; c) a histogram analysis has to be performed for each of the machine performance parameters (See Fig. 3). The distribution of parameters shows whether the performance parameters are rare and incidental (*p* < 0.05), and finally iv) features that contribute to optimal performance are extracted and compared to those chosen by the initial selection criteria. If those features are fully matched, we conclude that our feature selection is robust and reliable. In particular, for 1000 rounds, we a) randomly chose 37 vectors out of the 1792 features; b) calculated precision, sensitivity, specificity, and accuracy of the machine; d) saved the results in appropriate vectors and drew 4 histograms from 1000 element vectors of precision, sensitivity, specificity, and accuracy. Finally, as shown in Fig 3, the distribution of accuracy, precision, sensitivity and specificity show that our selected features are statistically, not random.

**Fig 3.**
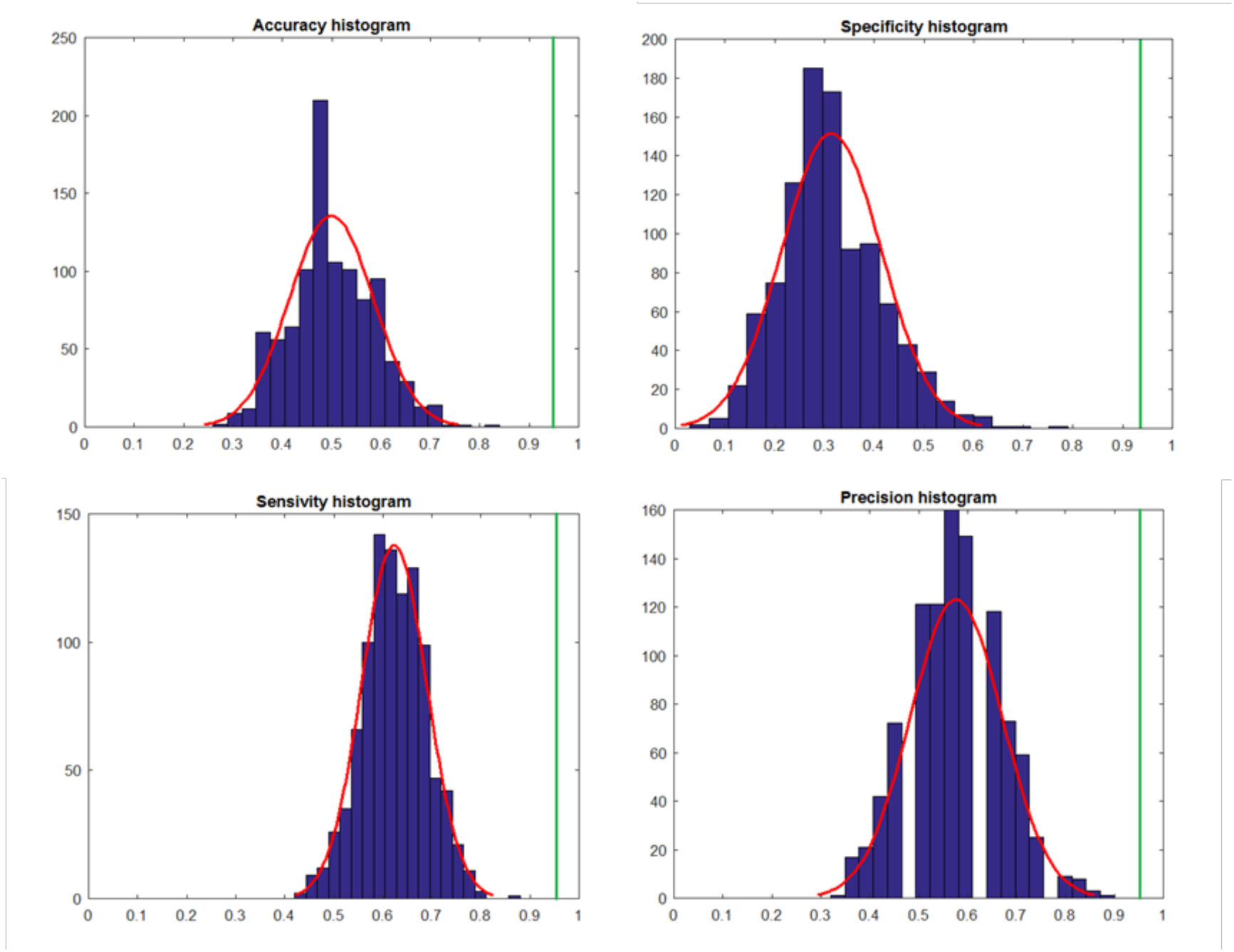
Random shuffling histogram. Red curve shows the distribution of accuracy, specificity, sensitivity and precision for 1000 rounds with the random shuffling of 37 features. The green line shows the results using the 37 FDR corrected features.

Our random shuffling results indicate that the selected features based on FDR corrected significant differences were robust and reliable.

## 5. Discussion

We applied two classifier to discriminate between dyslexics and typical readers based on local connectivity features derived from EEG resting-state. In our previous study, we showed group differences in how connectivity is organized between dyslexics and typical readers using graph measures that related to global properties of the network (Fraga González et al. 2016, 2018). In the current analysis, we focus on classification of subjects by using a larger number of features computed per node (i.e., scalp electrode). First, we found group differences associated with several local network measures that were selected to feed an automatic classifier. Then, our cross-validation analysis showed that the classifier could identify dyslexic readers with high accuracy (> 90%). Our random shuffling analysis showed that the selected features, most of which were associated with the delta frequency band, were useful to obtain a high accuracy classification that was not possible to achieve by a random selection of the network features. The current results are consistent with the data of a previous MEG study using sensor-level information (Dimitriadis et al. 2018). The results of this study showed that a higher classification of adults with reading difficulties was possible using sensor-specific network measures compared to using global (averaged across nodes) measures. The current results extend those and previous findings relating resting-state activity to reading abilities (Koyama et al. 2011) and suggesting differences in connectivity organization in dyslexia (Finn et al. 2014; Schurz et al. 2014).

Our classifier used network features derived from connectivity matrices in different frequency bands. The local network features describe different aspects of connectivity between the nodes, for example, how relevant a node is in the whole network or how well connected it is to its neighbors (e.g., Stam 2014). Importantly, oscillations at different frequency bands have different temporo-spatial features. Higher frequencies seem to relate to more local activity, i.e., smaller networks, while slower oscillations modulate larger networks with more widespread activity (Buzsáki and Draguhn 2004). It is likely that different aspects of dyslexia are manifested at multiple levels of neural organization. Indeed, previous research shows that reading impairments in dyslexia may result from a heterogeneous cluster of cognitive characteristics (Leinonen et al. 2001; Menghini et al. 2010; Pacheco et al. 2014). This is supported by evidence suggesting similarly complex and heterogeneous neurobiological profiles (e.g. Richlan et al. 2009; Finn et al. 2014). The current results suggest that EEG features provide important information about the underlying neurocognitive profiles in dyslexia, which can be used to classify individuals based on task-unspecific electrophysiological data. Recently, a study used structural and functional MRI data to classify a sample of 22 dyslexics and 27 typically reading adults (Tamboer et al. 2016). The study reported accuracy levels around 80% and detected specific occipital and parietal brain regions that contributed most reliably to the classification. We obtained high classification accuracy using local EEG features supporting the potential utility of using automatic classifiers in identifying individuals with dyslexia. However, the use of scalp EEG does not allow to draw conclusions about the underlying sources defining those features. Future studies using source modeling in MEG recordings could bring new insights into what sources are more relevant for classification.

For the present findings to have clinical implications, e.g., in diagnostics and early detection of reading disabilities, the generalizability of classification should be further investigated. In the study by Tamboer and colleagues (2016), classification performance dropped to around 60% when the trained classifier was applied to an independent sample. We only performed classification within the available dataset and it would be important to assess the current classifier’s performance within additional datasets (Pulini et al. 2018). In addition, to further advance on the relation between EEG network features and cognitive deficits in dyslexia, longitudinal studies should examine whether the current approach could be used to predict reading improvements or treatment outcomes. In relation to this, a previous study used structural and functional MRI data to predict future performance in children with dyslexia and found higher accuracy in prediction using brain measures compared to behavioral tests (Hoeft et al. 2007).

This study has several limitations. First, the current study included almost twice as many typical readers as readers with dyslexia. This bias is relevant to machine-learning based diagnostics as an imbalance sample results in a disproportionate representation of one of the classes in the training (Mazurowski et al. 2008). Secondly, this and other studies include a gap between typical readers (i.e., readers with a reading performance > 25%) and readers with dyslexia (i.e., readers with a reading performance < 10%). Obviously, such a gap is not present in the real world and might compromise our classifier when applied in a natural setting. Thirdly, the ratio of readers with dyslexia to typical readers is far off from the prevalence rates in the general population (1:20). Future studies should determine the sensitivity of the machine classifiers to changing odd ratios.

### General conclusions

This study builds upon the notion that dysfunctional connectivity between local specialized networks may be involved in dyslexia rather than global network measures studied in (Fragza et. al., 2016). We therefore focus on local network properties derived from EEG functional connectivity at rest and show that they can be used to classify individuals as dyslexics and typical readers. The current study presents an interesting and novel biologically-based method to analyze multidimensional data derived from EEG functional connectivity networks. Further research should help elucidating the clinical applicability of EEG-based classification and functional significance of these measures in relation to reading deficits in dyslexia. The current study adds to previous studies that encourage the application of machine learning techniques to neuroimaging data in order to improve subject classification in the context of reading disabilities (Tamboer et al. 2016; Dimitriadis et al. 2018). This approach may benefit from increasingly large data sets available for data-driven analysis.

## Acknowledgements

We would like to express our gratitude to all the children and parents for participating in the study.

## Supporting information

**S1 Table.**
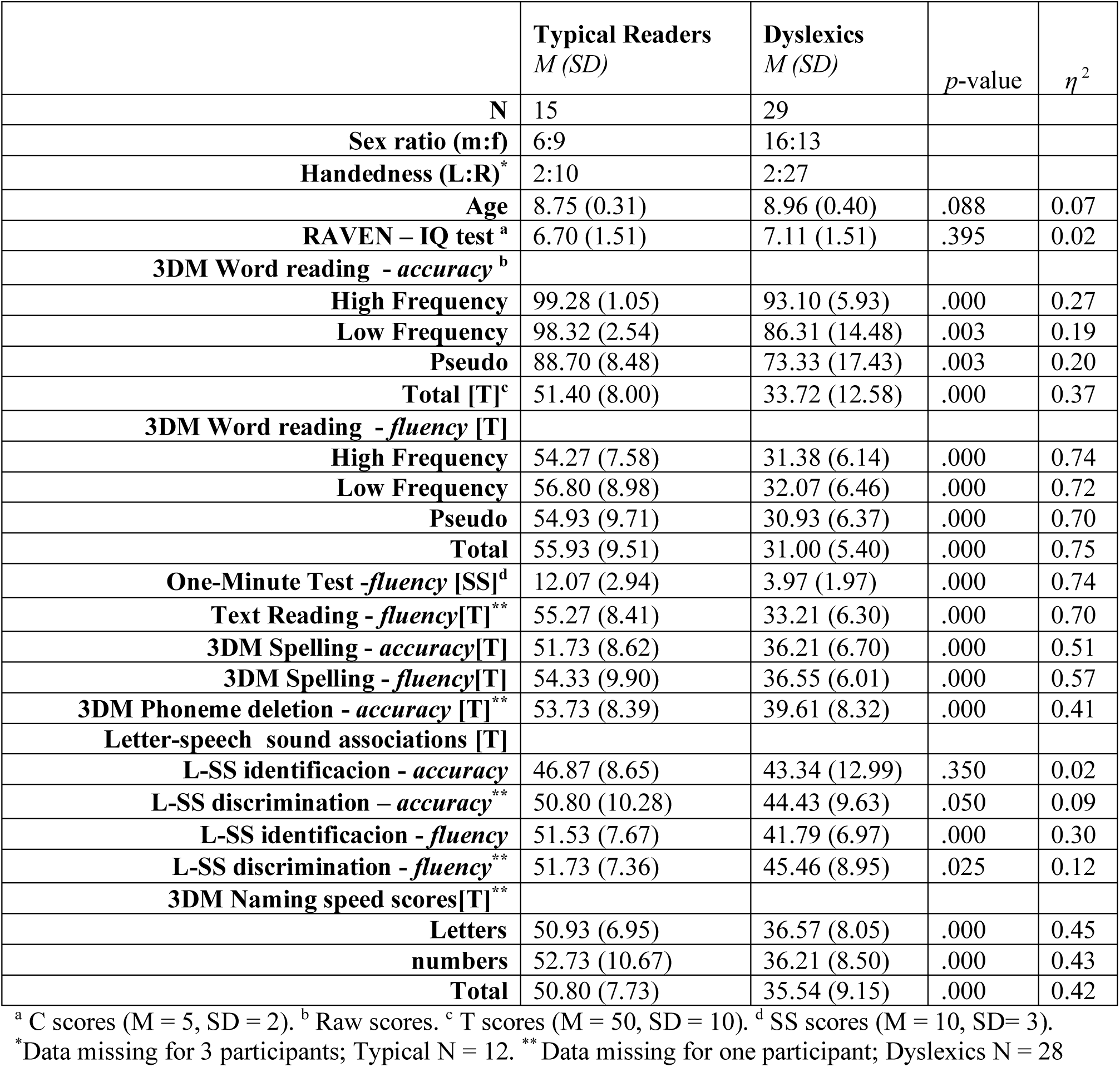
Sample characteristics and descriptive statistics of reading accuracy and fluency scores.

| | **Typical Readers**<br>*M (SD)* | **Dyslexics**<br>*M (SD)* | *p*-value | $\eta^2$ |
| --- | --- | --- | --- | --- |
| **N** | 15 | 29 |  |  |
| **Sex ratio (m:f)** | 6:9 | 16:13 |  |  |
| **Handedness (L:R)<sup>\*</sup>** | 2:10 | 2:27 |  |  |
| **Age** | 8.75 (0.31) | 8.96 (0.40) | .088 | 0.07 |
| **RAVEN – IQ test<sup>a</sup>** | 6.70 (1.51) | 7.11 (1.51) | .395 | 0.02 |
| **3DM Word reading - accuracy<sup>b</sup>** |  |  |  |  |
| **High Frequency** | 99.28 (1.05) | 93.10 (5.93) | .000 | 0.27 |
| **Low Frequency** | 98.32 (2.54) | 86.31 (14.48) | .003 | 0.19 |
| **Pseudo** | 88.70 (8.48) | 73.33 (17.43) | .003 | 0.20 |
| **Total [T]<sup>c</sup>** | 51.40 (8.00) | 33.72 (12.58) | .000 | 0.37 |
| **3DM Word reading - fluency [T]** |  |  |  |  |
| **High Frequency** | 54.27 (7.58) | 31.38 (6.14) | .000 | 0.74 |
| **Low Frequency** | 56.80 (8.98) | 32.07 (6.46) | .000 | 0.72 |
| **Pseudo** | 54.93 (9.71) | 30.93 (6.37) | .000 | 0.70 |
| **Total** | 55.93 (9.51) | 31.00 (5.40) | .000 | 0.75 |
| **One-Minute Test -fluency [SS]<sup>d</sup>** | 12.07 (2.94) | 3.97 (1.97) | .000 | 0.74 |
| **Text Reading - fluency[T]<sup>\*\*</sup>** | 55.27 (8.41) | 33.21 (6.30) | .000 | 0.70 |
| **3DM Spelling - accuracy[T]** | 51.73 (8.62) | 36.21 (6.70) | .000 | 0.51 |
| **3DM Spelling - fluency[T]** | 54.33 (9.90) | 36.55 (6.01) | .000 | 0.57 |
| **3DM Phoneme deletion - accuracy [T]<sup>\*\*</sup>** | 53.73 (8.39) | 39.61 (8.32) | .000 | 0.41 |
| **Letter-speech sound associations [T]** |  |  |  |  |
| **L-SS identificacion - accuracy** | 46.87 (8.65) | 43.34 (12.99) | .350 | 0.02 |
| **L-SS discrimination – accuracy<sup>\*\*</sup>** | 50.80 (10.28) | 44.43 (9.63) | .050 | 0.09 |
| **L-SS identificacion - fluency** | 51.53 (7.67) | 41.79 (6.97) | .000 | 0.30 |
| **L-SS discrimination - fluency<sup>\*\*</sup>** | 51.73 (7.36) | 45.46 (8.95) | .025 | 0.12 |
| **3DM Naming speed scores[T]<sup>\*\*</sup>** |  |  |  |  |
| **Letters** | 50.93 (6.95) | 36.57 (8.05) | .000 | 0.45 |
| **numbers** | 52.73 (10.67) | 36.21 (8.50) | .000 | 0.43 |
| **Total** | 50.80 (7.73) | 35.54 (9.15) | .000 | 0.42 |

<sup>a</sup> C scores (M = 5, SD = 2). <sup>b</sup> Raw scores. <sup>c</sup> T scores (M = 50, SD = 10). <sup>d</sup> SS scores (M = 10, SD= 3).
<sup>\*</sup>Data missing for 3 participants; Typical N = 12. <sup>\*\*</sup> Data missing for one participant; Dyslexics N = 28

